# Prenatal alcohol exposure alters axon formation and Robo/Slit guidance cue expression in the internal capsule at E15.5

**DOI:** 10.1101/2022.05.11.491520

**Authors:** Peter Chi, Kevyn Dewees, Jennifer Scoville, Terrence Deak, Carlita Favero

## Abstract

We previously showed the prenatal alcohol exposure (PAE) induced precocious growth of E15.5 internal capsule axons. We set out to further delineate which axon tracts were affected and a potential mechanism for this phenomenon. Pregnant Swiss Webster mice were exposed to saline or 20% ethanol via subcutaneous injection from embryonic days (E) 7.5 until E14.5. E15.5 embryos were captured via C-section and brains were either fixed for immunostaining or flash frozen for tissue punches. To identify internal capsule axons we performed immunohistochemistry using L1 antibody to visualize all internal capsule axons and NetrinG1 antibody to identify thalamocortical axons specifically. mRNA was extracted from tissue punches to visualize Slit1, Robo1, Slit2, and Robo2 gene expression levels via real time RT PCR. We found that L1-expressing axons exhibited increased axon crossing in females with the opposite effect in males and no difference in Netrin-G1-expressing axons. We also found PAE increased Robo1 mRNA. Taken together, our results indicate that PAE elicits specific effects on axon trajectories, potentially via the Slit/Robo guidance family of repellents and receptors. These findings add to the growing body of evidence demonstrating alcohol disruptions of axon growth and guidance during embryonic development that may contribute to connectivity errors that characterize Fetal Alcohol Spectrum Disorders.

## Description

We first set out to investigate whether internal capsule axons were altered along their trajectory in FASD mice. Internal capsule axons are comprised of thalamocortical axons, corticothalamic axons, and corticospinal axons which are involved in proper sensory and motor processing. FASD commonly results in sensory processing difficulties, so we hypothesized these axon tracts would be influenced by prenatal alcohol exposure (PAE)^1^. As a metric of axon growth, we investigated how PAE affected axon crossing at a critical checkpoint along their trajectory, the corticostriatal boundary (CSB). For this analysis, we assayed the number of L1-positive axons crossing the CSB at E15.5, a key developmental milestone along the trajectory^2^. L1 is a neural cell adhesion molecule expressed on all three axon tracts. We previously showed that L1-positive axons were longer at this age in PAE mice^3^, thus we wanted to investigate if PAE changed the number of axons crossing this checkpoint. Analysis revealed a significant Treatment X Sex interaction (t57 = 2.78, p = 0.007, Figure). We found opposing effects based on sex, with more L1-positive axons crossing in female FASD mice and less axons crossing in male FASD mice (Figure). Looking at high magnification images, it appears that the differences in crossing are due to extent of axon coverage, possibly relevant to size of axon fibers rather than change the number of axons. It is not uncommon to see males and females differentially affected from PAE. Literature consistently shows sex differences in social and anxiety behaviors^4^, auditory processing^5^, and postnatal microglial colonization^6^. We were particularly interested in thalamocortical axons because they carry sensory information from the thalamus to the cortex to be integrated (Reviewed in ^7^). To determine whether the differences we observed were due to thalamocortical axons, we assayed CSB crossing in netrin-G1-immunostained brain sections from E15.5 offspring. Netrin-G1 is lipid-anchored protein involved in thalamocortical axon guidance that is specifically expressed on these axons^8^. We did not find any differences in axon crossing in Netrin-G1 sections which presumes that the alcohol effects on L1 axon crossing were due to corticothalamic or corticospinal neurons.

**Figure 1.**
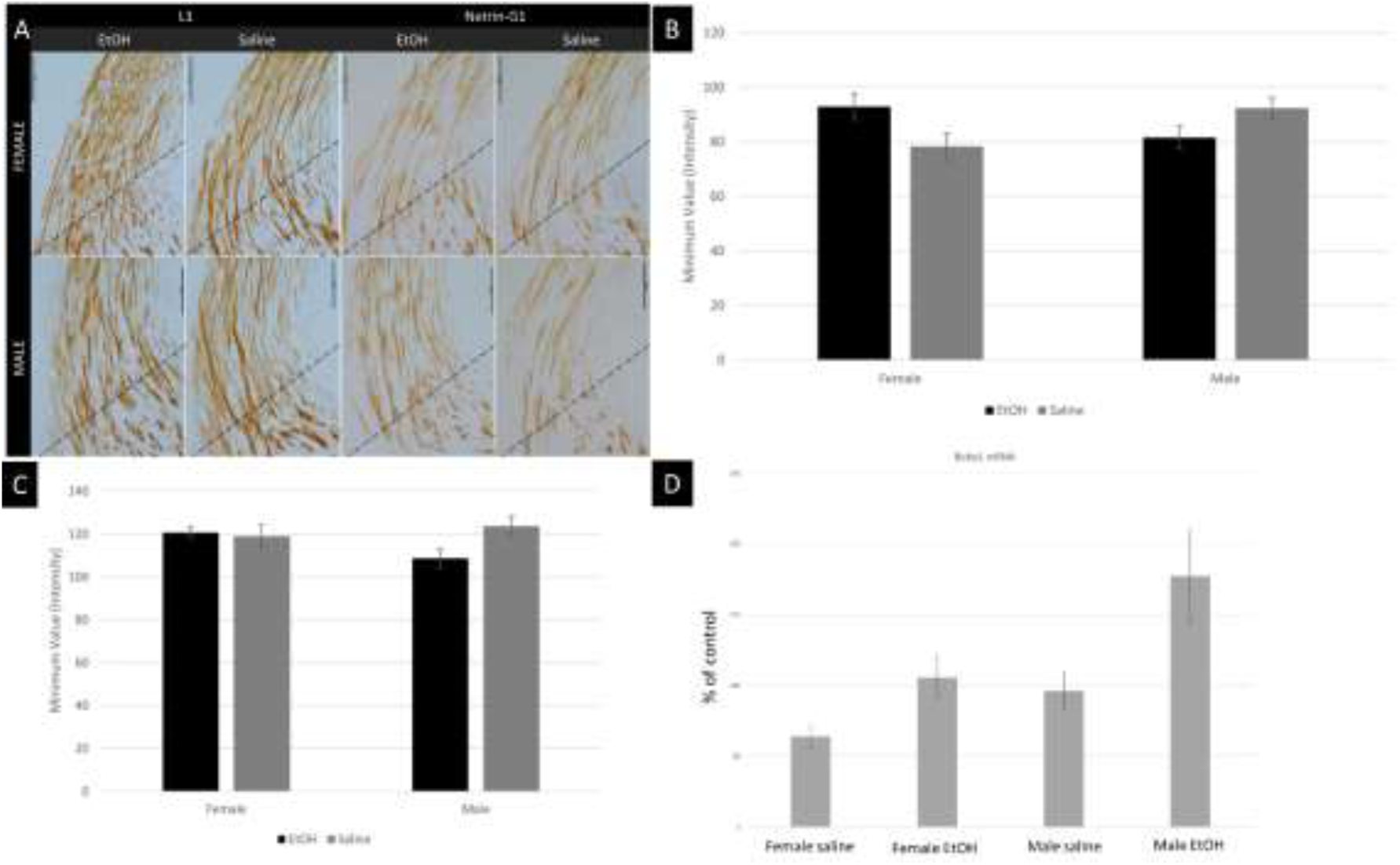
(A) Prenatal alcohol exposure (PAE) alters L1-expressing, but not Netrin-G1-expressing internal capsule axons. Representative images either show L1 or Netrin-G1 immunostaining on E15.5 coronal sections. L1 labels thalamocortical, corticothalamic, and corticospinal axons. Netrin-G1 labels thalamocortical axons only. Dotted lines represent the CSB, stretching from the sulcus to pial surface, from which a minimum value was calculated. Alcohol-exposed females had greater L1 staining intensity than saline-exposed controls. The opposite effect was found in males. There were no significant differences in NetrinG1 staining. Scalebar (vertical) = 100um. (B) PAE affects CSB crossing of L1-expressing axons, with differential effects in males and females. Lower min values correspond with higher staining intensity across the CSB. Alcohol significantly affected CSB crossing of L1-expressing axons (p = 0.041) with less crossing in females and more crossing in males. The interaction between sex and treatment was also significant (p = 0.007), with a trend towards a main effect of sex (p = 0.096). Data represent the mean. Error bars represent the standard error of the mean. (C) PAE does not affect CSB crossing of Netrin-G1-expressing axons. There was no main effect of treatment (p = 0.614) and no interaction (p = 0.155); similarly for L1, there was a trend towards a main effect of sex (p = 0.077). Data represent the mean. Error bars represent the standard deviation. (D) In alcohol-exposed embryos, we found a significant main effect of treatment for Robo1 mRNA expression in the internal capsule (t_22_ = -2.375, p=0.027) with highest magnitude levels in the alcohol injected group.

All internal capsule axons express Robo receptors and are guided by Slit repulsive ligands within this region^9^. Thus, we used real time RT PCR to measure mRNA levels of Robo1 and Slit 1 in the internal capsule of E12.5 embryos. Given our findings with altered length and crossing of internal capsule axons, we hypothesized that the aberrant growth was caused by altered Robo and/or Slit levels. E12.5 was chosen because this time point is just before these axons enter the internal capsule. Our hypothesis also supported findings from Vangipurum et al.^10^ showing that alcohol treatment of stem cell cultures reduced Robo1 mRNA. In alcohol-exposed offspring, we found a significant main effect of treatment for Robo1 (t_22_ = -2.375, p=0.027) with highest magnitude levels in the alcohol injected group and no difference in Robo2, Slit1, or Slit2 (data not shown) mRNA expression. One might presume that increased Robo1 expression would further repel internal capsule axons resulting in reduced outgrowth, but that doesn’t match our findings. Alternatively, Robo1 expression could be increased as a compensatory mechanism for malfunctioning Robo receptors. Since we did not test Robo1 function, we cannot rule out this possibility.

There is very little research into effects of PAE on growth and guidance of axon populations (our current understanding reviewed in ^11^, but several studies provide evidence these processes may be affected. Minciacchi et al.^12^ show that the cortical zone that receives thalamocortical axons is thinner in ethanol-exposed rats, and that thalamo-cortical termination is abnormal in the cortex. Granato et al.^13^ used thalamic and cortical injections of lectin-conjugated horseradish peroxidase to visualize axons, and they found that axon terminal fields for both TCA and CTA projections were altered in alcohol-exposed, most often reducing arborization. Thus, there are many potential areas of alcohol impact on internal capsule axons. Here we have looked at guidance and growth at a critical decision point, but other aspects of growth, such as initiation or termination, could be affected in place of or along with these. It is important that we continue this research to understand the specific mechanisms affected by PAE and aid in developing therapies for behavioral and cognitive deficits present in FASD.

## Methods

### Mice and treatment administration

All experiments were conducted using methods approved by the Institutional Animal Care and Use Committee of Ursinus College and in accordance with the National Institute of Health Guide for Care of Laboratory Animals. Experimental subjects were offspring of timed-pregnant Swiss Webster mice purchased from Charles River Laboratories (Raleigh, NC). Dams were delivered at gestational day E2.5 and housed in groups of three or four in polycarbonate cages with paper bedding on a 12-h reverse light-dark cycle. Food (Rodent Diet 5001, LabDiet) and tap water were freely available. Pregnant dams were randomly assigned to receive daily subcutaneous injections of 15uL per gram of bodyweight (2.3 g/kg) of either 0.9% saline or 20% (v/v) ethanol (EtOH) diluted in saline. Injections began on E7.5 and continued through E14.5. This dose results in maternal plasma blood alcohol concentration (BAC) of 194.7 +/-15.36 (mean +/-standard deviation) on E13.5 (n=3 dams) as determined using the Analox AM1 system (Analox Instruments USA, Lunenburg MA). To minimize stress of injection, dams were anesthetized using Isoflurane and injections were switched to the opposite side starting E11.5.

### Sex genotyping

Tails were removed from each embryo for DNA isolation. To distinguish between males and females, the Jarid1C and Jarid1D genes were amplified^14^. DNA extraction was performed using the Epoch Life Sciences *GenCatch* Genomic DNA Extraction kit (Epoch 2460050-1/13). We followed an adjusted version of the kit’s protocol. The attached protocol suggests using 200µL of heated water or Tris as an elution buffer, but we used 20µL of heated elution buffer (Buffer AE) from the Qiagen DNeasy Blood & Tissue Kit to maximize DNA yield (Qiagen 69506). Once DNA was isolated, concentrations were measured using a Nanodrop 2000 (Thermo Fisher). We prepared a stock solution of sex primers: 1µL of forward primer (5’-CTGAAGCTTTTGGCTTTGAG-3’), 1µL of reverse primer (5’-CCACTGCCAAATTCTTTGG-3’), and 100µL of sterile H_2_O. For each sample, we prepared this mixture for PCR: 10µL of Taq polymerase (Sigma P0982), 1µL of the premade sex primer stock, 200ng of DNA, and enough sterile H_2_O to fill the volume to 20µL. Samples ran through a PCR program of 35 cycles in the following: denaturing for 20 seconds at 94°C, elongation for 60 seconds at 54°C, and extension for 40 seconds at 72°C. Following PCR, the samples were run on a 2% agarose gel diluted in 0.5X TBE at 100V for approximately 90 minutes.

### Immunohistochemistry

To visualize internal capsule and thalamocortical axons in the embryonic brain, we stained E15.5 coronal brain sections with L1 and Netrin-G1, respectively. Slides were chosen that included the internal capsule. The immunostaining procedure started with slide dewaxing and rehydration. This consisted of slide washes in the following order: Histoclear (5 min, Electron Microscopy Sciences #64110), 100% EtOH (5 min), 95% EtOH (2 min), 70% EtOH (2 min), DH_2_O (2 min). This was followed by a 20-minute microwave incubation in 1% antigen retrieval citric acid-based solution (Vector H-3300 in DH_2_O). After cooling, we rinsed slides in 3 changes of 1 x phosphate-buffered saline (PBS) for 5 minutes each. Each slide was then treated with 200µl of the buffer solution (10% normal goat serum and 0.1% triton in 1 x PBS). After covering with parafilm, these slides sat for 30 minutes in a humidified chamber. Three 10-minute washes in PBS + 0.05% Tween followed this incubation. We then treated each slide with 200µl of the anti-L1 antibody (AbCam #ab208155, 1:250 in 1 x PBS) or anti-Netrin-G1 antibody (R&D Systems #AF1166, 1:100 in 1 x PBS). Parafilm covered these slides overnight in a humidified chamber at 4ºC. The following day, all slides went through three washes of PBS + 0.05% Tween for 10 minutes each, followed by a 30-minute incubation after treatment with ImmPRESS goat anti-rabbit secondary antibody (Vector #MP-7401) for L1 stained slides or ImmPRESS horse anti-goat secondary (Vector #ZH-0526) for Netrin-G1 stained slides. Parafilm covered the slides during incubation and was subsequently removed after the 30 minutes and slides were put through the following rinses: 10 minutes in PBS + 0.1% Tween, 2 × 5 minutes in 1 x PBS, 5 minutes in DH_2_O. We then used diaminobenzidine (DAB, Vector #SK-4105) to visualize the stain. Each slide got 200µl of DAB solution and sat for 1-2 (L1) or 5 (NetrinG1) minutes before being placed in DH_2_O. Finally, slides were dehydrated – 2 min in 70% EtOH, 2 min in 95% EtOH, 5 min in 100% EtOH, 5 min in Histoclear – and left to dry before coverslipping with Cytoseal.

### Imaging and CSB Crossing Analysis Protocol

We used a Nikon Eclipse 80i Compound Microscope with Nikon C2-SH digital camera to visualize staining at 4x magnification for L1- and Netrin-G1-stained slides. We also imaged TH slides at 10x for better visualization. We captured images of these stained slides using NIS-Elements AR software. To measure axon density along the corticostriatal boundary (CSB), we utilized ImageJ software to measure the intensity of pixels along a line. Lines were drawn using the straight-line tool from the sulcus^15^ to the pial surface of the brain (line angle = -150º for left hemisphere, -30º for right hemisphere). A second line was drawn directly above and parallel to the first line, covering only the area containing visibly stained axons. “Analyze” > “Measure” provided line length, average values, minimum values, and maximum values, all of which we recorded for each analyzed section. Minimum values correspond with staining intensity. Statistical analysis was only performed on minimum values. We remained blind to treatment, genotype, and sex during the time of analysis.

### Real time RT PCR

Brain tissue was harvested using rapid, unanesthetized decapitation. Heads were stored at -80C until subsequent tissue dissection using a core-sampling punch technique (as described in ^16^) for the extraction of the internal capsule. Gene expression in the brain was analyzed using real time reverse transcription polymerase chain reaction (RT-PCR), as described elsewhere^17^. Briefly, tissue was placed into a 2.0ml Eppendorf tube containing 500uL Trizol RNA reagent and a 5mm stainless steel bead, and then homogenized using a Qiagen Tissue Lyser II. Following homogenization, RNA was extracted using Qiagen’s RNeasy mini kit (catalog #74106), according to manufacturer’s instructions. Synthesis and storage of cDNA included a DNAse treatment step, and probed cDNA amplification was performed and captured in real time using the CFX384 Real-Time PCR Detection System (Bio-Rad, #185-5485). All PCR data were adjusted using the 2^-ΔΔC(T)^ method^18^ and are shown relative to a stable reference gene and expressed as percent of control. Glyceraldehyde 3-phosphate dehydrogenase (GAPDH) was used as the housekeeping reference gene.

## Statistical Analysis

All analyses proceeded in the linear model framework, for each outcome of interest. Due to the fact that different effects in each sex are expected, we not only included sex as a covariate in each model, but also a treatment x sex interaction term. For axon crossing analyses, due to some repeated measurements on each individual, we utilized a linear mixed model approach with a random effect for mouse ID, to account for correlation within individual mice as appropriate. All analyses were carried out in the R Statistical Programming Language^19^, and the package nlme^20^.

## Acknowledgements

We would like to thank Hira Khattak, Katie Renzi, Betty Endeshaw, and Ixchel Mendez for being excellent labmates and thoughtfully reflecting on this data.

